# Chemically Induced Chromosomal Interaction (CICI): A New Tool to Study Chromosome Dynamics and Its Biological Roles

**DOI:** 10.1101/2020.01.01.892448

**Authors:** Manyu Du, Fan Zou, Yujie Yan, Lu Bai

**Affiliations:** Department of Biochemistry and Molecular Biology, The Pennsylvania State University, University Park, PA, 16802; Center for Eukaryotic Gene Regulation, The Pennsylvania State University, University Park, PA, 16802; Department of Physics, The Pennsylvania State University, University Park, PA, 16802

## Abstract

Numerous intra- and inter-chromosomal contacts have been mapped in eukaryotic genomes, but it remains challenging to link these 3D structures to their regulatory functions. To establish the causal relationships between chromosome conformation and genome functions, we need a method that allows us to selectively perturb the conformation at targeted loci. Here, we developed a method in budding yeast, Chemically Induced Chromosomal Interaction (CICI), to engineer long-distance chromosomal interactions selectively and dynamically. We implemented CICI at multiple intra- and inter-chromosomal loci pairs and showed that CICI can form in >50% of cells, even between loci with very low Hi-C contact frequencies. CICI formation is slower at these low Hi-C sites, revealing the dynamic nature of the Hi-C signals. As a functional test, we forced the interaction between mating-type locus (*MAT*) and *HMR* and observed significant change in donor preference during mating-type switching, showing that chromosome conformation plays an important role in homology-directed DNA repair. Overall, these results demonstrate that CICI is a powerful tool to study chromosome dynamics and the 3D genome function.

---

3D chromosome conformation is thought to play critical roles in various nuclear processes, including DNA replication ^1^, repair ^2-4^, and transcription regulation ^5-9^. Recent applications of the Chromosome Conformation Capture techniques (3C, 4C, Hi-C, etc.) have discovered numerous intra- and inter-chromosomal interactions ^10-13^. We currently have a limited understanding of the function of these long-distance chromosomal interactions. Many studies have examined the *correlation* between chromosome organization and genomic processes, but it remains challenging to investigate their *causal* relationships.

To illustrate causality, the traditional genetic approach is to identify a protein that establishes the chromosomal interaction, disrupt the protein, and probe the biological consequences. Unfortunately, some of these proteins, like cohesin and CTCF, are essential for maintaining the overall 3D structure of the chromosomes, and mutations of these proteins will generate global and pleiotropic effects. In addition, proteins that set up chromosome interactions may also directly participate in genomic processes. For example, the looping between the Locus Control Region (LCR) enhancer and the β-globin promoter is established by transcription factors GATA1, FOG1, KLF1 and Ldb1 ^14-16^; the clustering of *GAL* genes or heat-shock genes requires transcription factor Gal4 or Hsf1 ^17,18^. Mutations of these factors will result in simultaneous loss of chromosomal interactions and gene expression, preventing us from elucidating the relation between them.

We therefore need a method to selectively perturb chromosomal interactions without directly disrupting nuclear functions. A number of solutions have been presented in recent years, including artificial tethering of chromosomes through tetrameric LacI ^19,20^, forced chromosome looping by tethering dimerization factors with zinc-finger proteins ^21^ or dCas9 ^22,23^. However, it is not clear what fraction of cells can form forced interactions with these methods, and their applications so far have been largely limited to well-studied enhancer-promoter systems like the LCR-β-globin region. Some of these methods rely on constitutive protein-protein interactions, and therefore the interactions they generate are not dynamically controlled. Here, we 1) developed a method that can rapidly induce chromosomal interactions between different pairs of targeted loci, 2) investigated the relations between the rate / efficiency of the induced interaction and the Hi-C contact frequencies of these loci, and 3) perturbed the chromosome conformation near the mating locus and probed the consequences in DNA repair.

The method we developed, Chemically Induced Chromosomal Interactions (CICI), involves two existing techniques, chemically induced dimerization (CID) ^24,25^ and fluorescent repressor operator systems (FROS) ^26,27^. In a commonly-used CID system, FK506 binding protein (FKBP12) and the FKBP-rapamycin binding domain (FRB), two otherwise non-interacting factors, bind strongly in the presence of rapamycin ^28^. We expressed LacI-FKBP12 and TetR-FRB fusion proteins in a rapamycin-resistant yeast strain containing LacO and TetO repeats at desired genomic locations (**Fig 1A**). In addition to the FKBP12 and FRB fusions, we also introduced LacI-GFP and TetR-mCherry into the same strain for visualization. We expected the LacI-FKBP12 and TetR-FRB to bind tightly with each other in the presence of rapamycin, resulting in the co-localization of the LacO and TetO “chromosome dots” (**Fig 1A**). This experimental scheme is based on exogenous factors that have no specific binding partners in the native genome, so the interactions should be established with high specificity at the target sites. Large number of FKBP12-FRB contacts over the LacO and TetO repeats (256X and 192X) should stabilize the chromosomal interactions. Addition of rapamycin allows the interaction to be dynamically induced.

**Fig 1.**
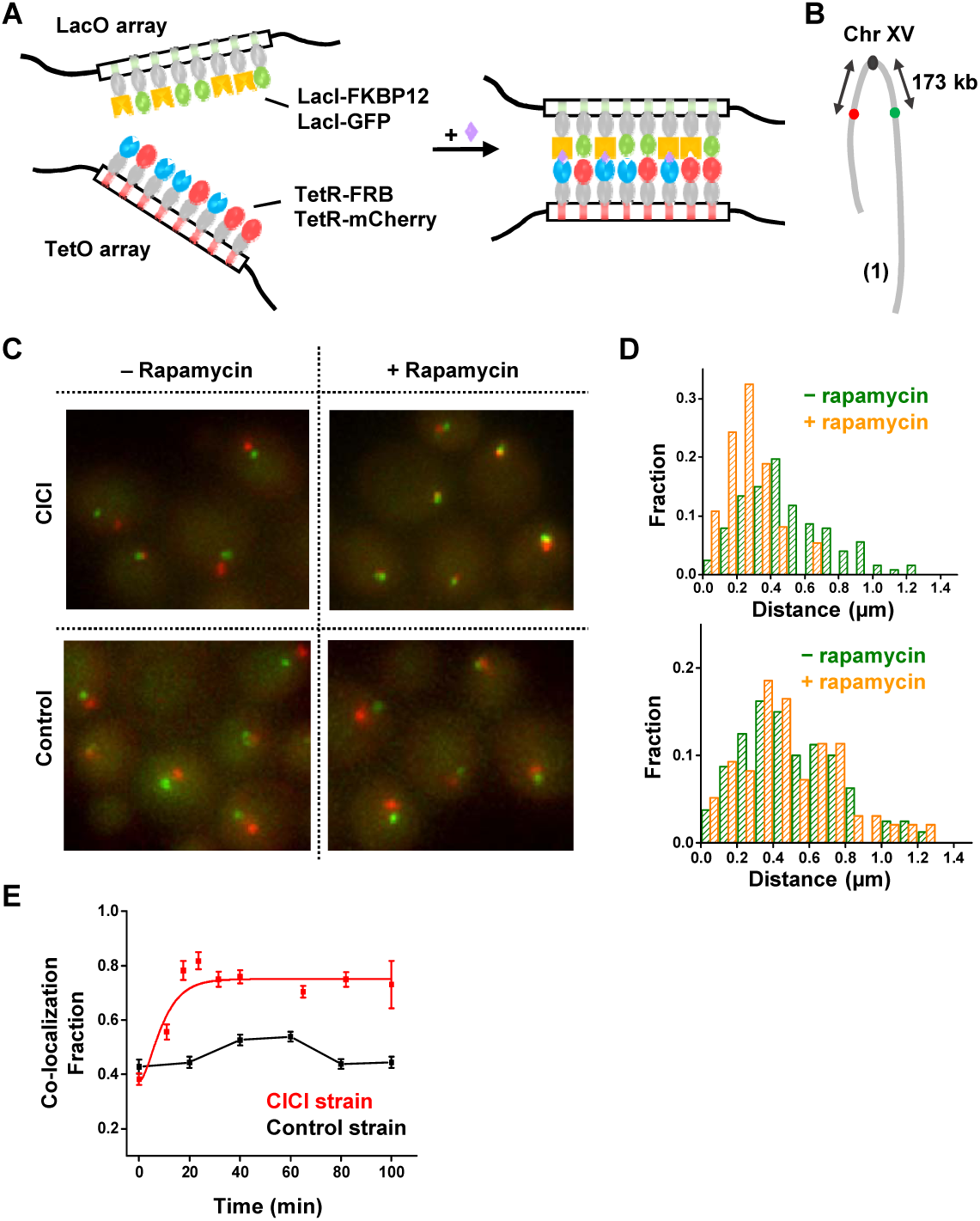
Implementation of CICI. **A)** The scheme of CICI. LacO and TetO arrays were inserted into two genomic loci. Fusion proteins LacI-FKBP12 and TetR-FRB associated with these arrays dimerize in the presence of rapamycin. LacI-GFP and TetR-mCherry were also included for visualization. **B)** LacO and TetO arrays are integrated 150 kb from the centromere on the left and right arms of Chr XV. **C)** Visualization of the CICI strain (containing the complete CICI system in (A)) and the control strain (containing LacI and TetR instead of LacI-FKBP12 and TetR-FRB) ± rapamycin. **D)** Histograms of distances between the two dots in the CICI strain (upper panel) and in the control strain (lower panel) ± rapamycin. In total, 128, 95, 80, and 97 cells were measured respectively. **E)** Fraction of co-localization as a function of rapamycin incubation time in the CICI (red) or the control strain (black). Total # of data points were 358 for CICI and 892 for control.

For this assay to work, it is essential to keep the concentrations of the LacI-FKBP12 and TetR-FRB fusion proteins at low levels: freely diffusing proteins dimerize with the chromatin-bound counterparts and therefore interfere with the interaction between the two targeted loci. Based on our previous study ^29^, we constructed strains containing the *REV1* promoter driving LacI and TetR fusion proteins, which allows high occupancy of LacO / TetO arrays with low fusion protein concentrations (**Methods**).

As a proof of principle, we first tested CICI with a loci pair that is known to have a high tendency to interact. Chromosomal interactions in yeast are strongly constrained by the Rabl configuration where centromeres cluster, and the neighboring chromosome arms run closely in parallel ^30-32^. As a result, Hi-C experiments showed that loci with equal distances to centromeres have higher contact frequencies ^31^. We therefore inserted LacO and TetO arrays into the left and right arm of Chr XV, both ∼ 173 kb away from the centromere (pair 1) (**Fig 1B**). We incubated these cells with or without rapamycin for 1 hr, imaged under a fluorescent microscope, and analyzed the distance between the LacO and TetO dots in individual cells (**Methods**). The LacO and TetO dots become closer in the presence of rapamycin (average distance decreases from 0.51 µm to 0.26 µm) (**Fig 1C & D**). In contrast, LacO-TetO distance in a control strain that lacks the CID components (LacI and TetR instead of LacI-FKBP12 and TetR-FRB) does not change with rapamycin (**Fig 1C & D**), showing that the LacO-TetO co-localization is not a generic reaction to rapamycin, but represents true CICI events. To measure the rate of CICI formation, we imaged cells at different time points after adding rapamycin. The co-localization probability rapidly increases from ∼40% to ∼80% within 20 min in the CICI strains, but remains constant in the control strains (**Fig 1E**).

Given that CICI can be successfully established for the loci above, we next generated four more strains with different LacO / TetO insertion sites (pair 2-5 in **Fig 2A**). These loci pairs were chosen strategically to span a broad range of contact frequencies based on published Hi-C data ^30^ (**Fig 2B**). In comparison to pair 1, the Hi-C signal is higher for pair 2 (symmetric to centromere with shorter linear distance) and lower for pair 3 (similar linear distance but asymmetric to centromere). The Hi-C signals for pair 4 and 5 are even lower. For pair 4, LacO and TetO were placed on the two sides of rDNA, which separates Chr XII into two isolated domains ^31^. For pair 5, LacO and TetO were inserted into two different chromosomes, Chr XV and Chr XII, and the frequencies of “inter-chromosomal” interactions are generally lower than those of “intra-chromosomal” interactions. Upon induction with rapamycin, we observed CICI formation in all of these strains (**Fig 2C**) despite the fact that some of the loci pairs are initially well-separated in the nucleus.

**Fig 2.**
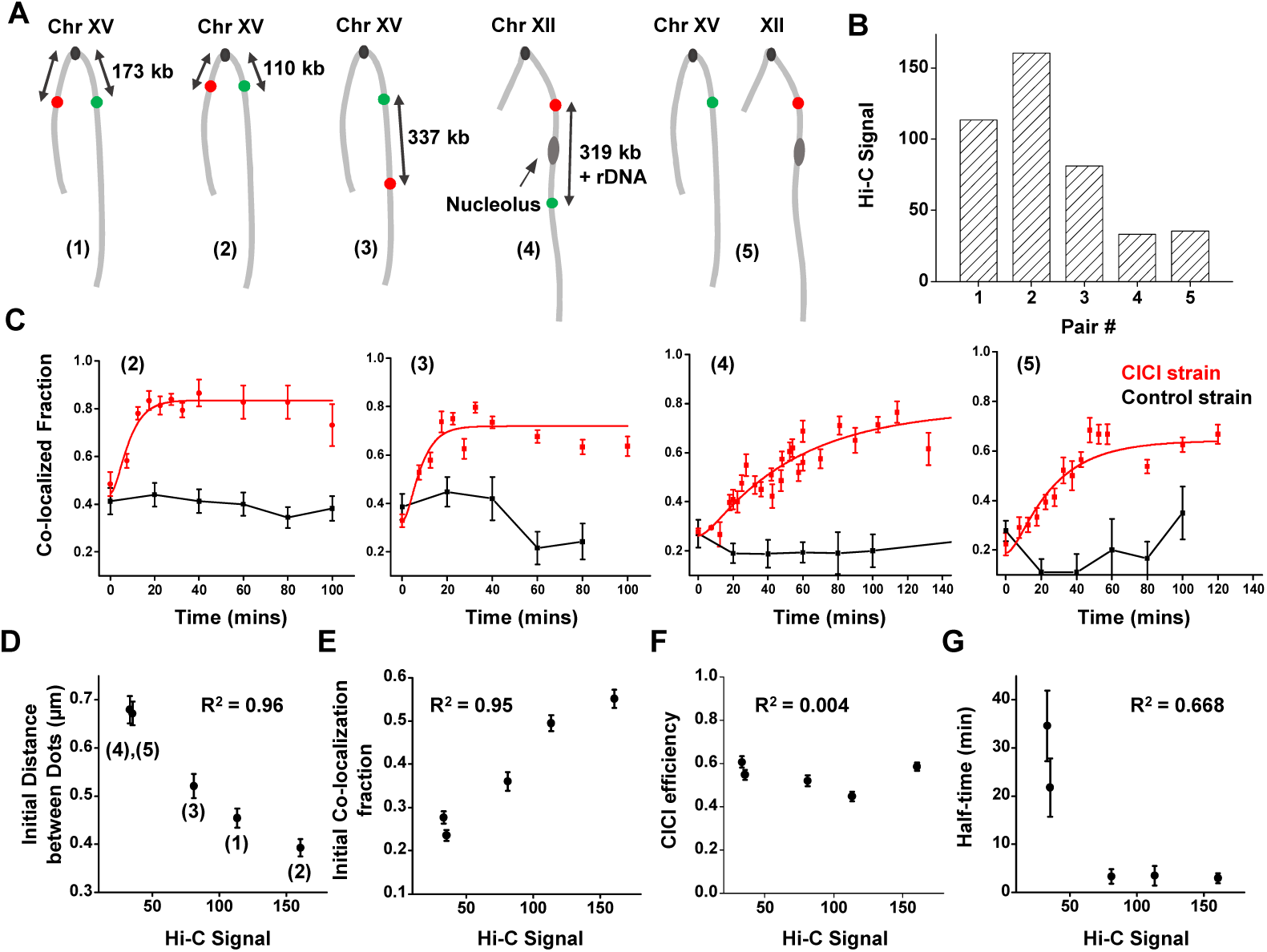
CICI formation at loci with different Hi-C contact frequencies. **A)** Five loci pairs chosen for the CICI test. **B)** Hi-C signals of the five pairs of loci measured previously ^30^. **C)** The dynamics of CICI formation for loci pair 2-5 (from left to right). Total # of data points were 321 (CICI) & 585 (control) for (2), 352 & 587 for (3), 399 & 311 for (4), and 168 & 114 for (5). **D-G)** The correlation between Hi-C signal with the initial distance between the dots (D), initial co-localization probability (E), CICI efficiency (F), and the half-time of CICI formation (G). Points represent loci pairs 4, 5, 3, 1, & 2 from left to the right. The corresponding coefficients of determination R^2^ are shown in the panels.

Using the CICI formation data above, we derived the initial mean distance between dots, initial co-localization probability, CICI efficiency (net co-localization gains at steady state), and *t*_½_ of CICI formation for each loci pair (**Methods**), and plotted these variables against their Hi-C signals. In the absence of rapamycin, the Hi-C signals show strong negative correlation with the dot distance, and strong positive correlation with the co-localization probability (R^2^ >0.95; **Fig 2D & E**). In particular, pair 4 and 5 have similar Hi-C signals and distances, even though that one is intra- and the other is inter-chromosomal. Such correlations are expected because Hi-C measures the probability for two loci to fall within a ligatable distance, which is largely determined by the average distance in-between. In striking contrast, the Hi-C signals have essentially no correlation with the CICI efficiency (R^2^ <0.01; **Fig 2F**), i.e. for all the CICI pairs generated, interactions can form in 50-60% of cells regardless of their initial 3D conformation. The rate of contact formation, however, depends on the initial conformation: for pair 4 and 5 with low Hi-C contact frequencies, CICI formation is almost one order of magnitude slower than the other three pairs (**Fig 2G**). Overall, these data reflect the *kinetic nature* of the Hi-C signals: two loci with low contact frequency can still form interactions, but it takes longer for them to encounter. This conclusion is consistent with analyses of chromosome conformation in higher eukaryotes, which revealed that loci pairs in different Topologically Associated Domains (TADs) can be in physical proximity in a fraction of cells ^33^. Based on our observation, we suspect that intra-TAD interactions have kinetic advantages over inter-TAD ones. However, interactions can eventually form when they are induced across TAD boundaries, especially in the absence of competition.

The ultimate goal of developing CICI is to perturb chromosome conformation and probe its biological consequences. Here, we examined the effect of CICI on DNA homology-directed repair. Long-range interactions between homologous sequences should play an essential role in this process as the repair template needs to be in close vicinity of the DNA lesion ^34,35^. Mating-type switching is a well-characterized model for homologous repair in budding yeast ^36,37^. The switching is initiated by the *HO*-mediated double-stranded break at the *MAT* locus, which is then repaired by *HMLα* or *HMRa* located on the opposite ends of Chr III. The donor preference (*MATa* is mostly repaired by *HMLα*, and *MATα* by *HMRa*) is thought to result from differential configurations of Chr III in a and α cells. However, *MAT-HML* and *MAT-HMR* contact frequencies only have mild differences in cycling a and α cells ^30^. Previous studies have used tetrameric LacI to alter the Chr III conformations ^20^ or membrane-tethered LacI to constrain the motion of *HML* ^38^, and observed some changes in the donor preference. However, how efficiently these methods perturb the proximity of *MAT-HML* and *MAT-HMR* is not clear. Here, we directly induced interaction between *MAT* and *HMR* in *MATa* (a mating type) cells to see if we could rewire the donor preference.

We constructed a *MATa* yeast strain containing the CICI fusion proteins with LacO and TetO ∼5 kb downstream the *MAT* and *HMR* loci (**Fig 3A**). We confirmed that CICI can form in this strain with a ∼50% efficiency (**Fig 3B & C**). This strain also has *GAL10pr-HO* and *HMRα-B* (*HMRa* replaced by *HMRα* with an engineered BamHI cutting site ^38^), allowing us to induce the *MAT* repair with galactose and probe the relative use of the *HML* and *HMR* based on BamHI digestion (**Methods**). We also generated the control strain that lacks FKBP12 and FRB. We grew these strains to log-phase in raffinose, added rapamycin for 1 hr to establish CICI, and induced the *HO* expression with galactose for 1 hr. We then added glucose to repress *HO* for 1 hr, allowing repairs to take place, before extracting the genomic DNA and genotyping the mating locus (**Methods**). The presence of rapamycin increases the usage of *HMRα-B* in the CICI strain from 6% to 27%, but shows no effect in the control strain (**Fig 3D & E**). Given that the CICI induces co-localization in 50% of cells, the *HMR* usage with the proximal *MAT-HMR* conformation should be higher than the population average (between 40-50%). These data demonstrate that the engineered chromosomal interaction in **Fig 3A** can significantly override the endogenous mechanism of donor selection during mating-type switching.

**Fig 3.**
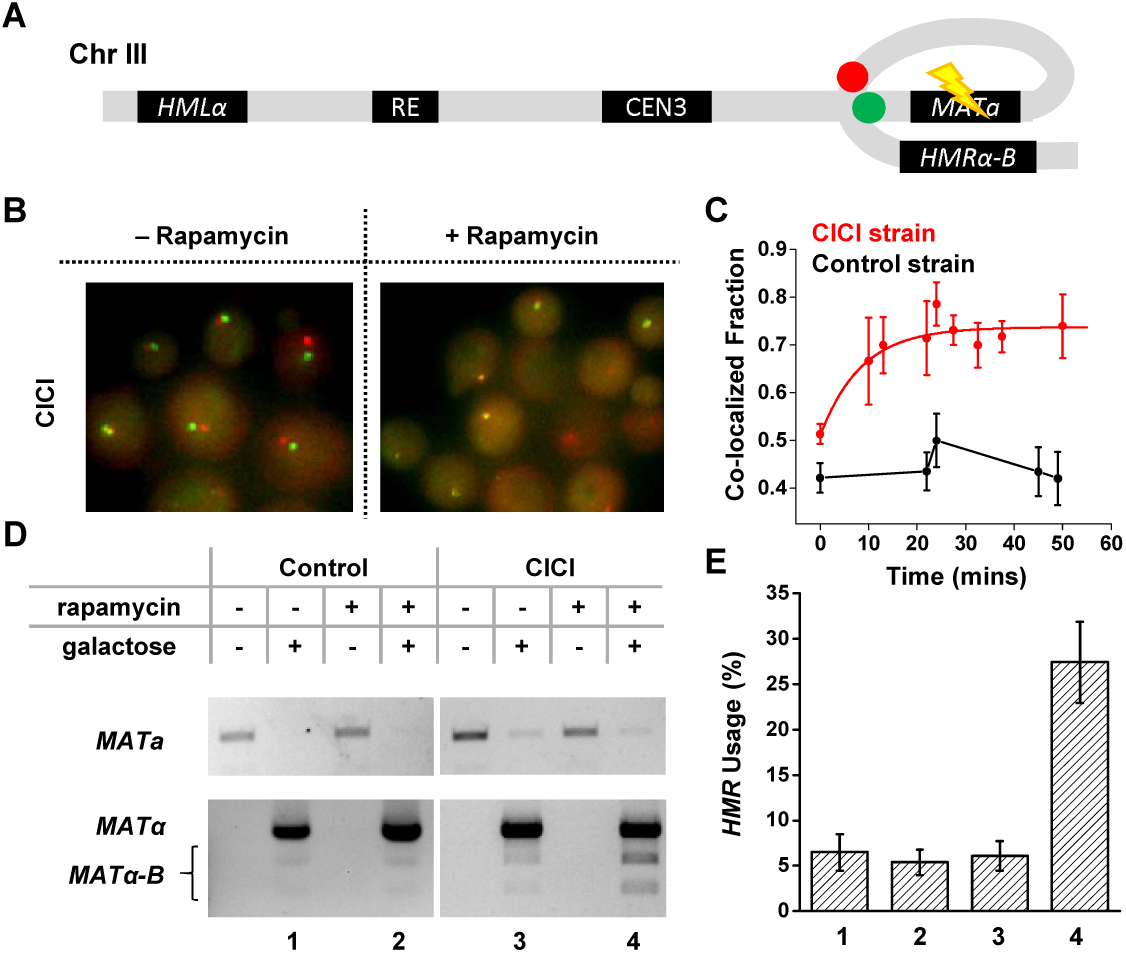
CICI changes the donor preference in homology-directed DNA repair. **A)** Engineered proximity between *MATa* and *HMRα-B* by CICI. Green and Red dots: LacO and TetO arrays; Flash symbol: HO digestion. **B)** Visualization of the *MATa* and *HMRα-B* loci ± rapamycin. **C)** Fraction of co-localization as a function of rapamycin incubation time in the CICI (red) or the control strain (black) used in the mating-type switching assay. Total # of data points were 226 for the CICI strain and 272 for the control. **D)** Quantitative analysis of *HMR* usage in yeast mating-type switching. Top gel: PCR of the *MATa* sequence, which decreases after the *HO* induction (representing the switching from a to α or α-B). Bottom gel: PCR of the *MATα* sequence followed by BamHI digestion. Undigested / digested bands represent the usage of *HML* (α) and *HMR* (α-B). **E)** The fraction of *HMR* usage for the four conditions in D. Standard error was calculated from two or three biological repeats.

In this study, we developed a synthetic biology method, CICI, to induce chromosomal interactions between targeted loci. Using this method, intra- and inter-chromosomal interactions can be engineered and dynamically controlled through a diffusive chemical. CICI can form between loci with very low Hi-C contact frequencies in a large fraction of cells, although the formation takes longer. Such CICI formation time reflects the rate of chromosome encounter, which provides insight into chromosome dynamics. Combining CICI with a DNA repair assay revealed that perturbation in chromosome conformation can alter the donor choice of homologous recombination. In doing so, this study goes beyond correlation and directly establishes the causal function of chromosome topology in this process. In yeast, chromosome conformation was also thought to affect transcription ^17,39-42^ and DNA replication ^43^. In the future, CICI can be used to study these 3D genome functions. Because both CID and FROS have been implemented in multicellular eukaryotic cells ^44,45^, we expect that CICI can be constructed in these model organisms and have broad applications.

## Methods

### Plasmid and strain construction

Standard methods were used to construct the strains and plasmids (see **Table S1** for plasmid and strain information). All the strains used in this study were derived from a background strain carrying *tor1* and *fpr* mutations for rapamycin resistance. The insertion loci were selected to be within intergenic regions between convergent gene pairs to have minimal effects on the endogenous gene expression (see **Table S1** for insertion index information). We constructed the CICI strain by integrating the plasmid containing LacI-FKBP12 and TetR-FRB driven by *REV1pr* into the *ADE2* locus. For the control strain, we integrated the plasmid containing LacI and TetR driven by *REV1pr*. We then integrated a plasmid containing *REV1pr*-LacI-GFP and *REV1pr*-TetR-mCherry into the *HIS3* locus to allow fluorescence detection of the LacO and TetO arrays. Since these arrays contain repetitive sequences that are difficult for molecular cloning, we used a two-step integration method developed by the Gasser lab ^46^. We first integrated an *URA3* selection marker into the target site, and then replaced *URA3* by LacO array through FOA selection. The same protocol was repeated subsequently for the TetO integration, but with a different sequence flanking the *URA3* marker. After every step of strain construction, yeast colony PCR was performed to verify the integration site. For strains used in the mating-type switching assay, we integrated the LacO/TetO array into the intergenic regions ∼ 5 kb downstream the *MAT* and *HMR* loci with the same method, so that the insertion would not disrupt the mating loci, and the homologous recombination could happen in the right orientation (see **Table S1** for insertion index information). We also mutated the *HMRa* sequence to *HMRα-B*, which can be differentiated from *HMLα* by by BamHI digestion. This manipulation is done through mating with a strain containing *HMRα-B* (generously provided by Dr. Haber) and tetrad dissection. Additionally, we inserted a *GAL10pr-HO* into the *GAL10* locus to express *HO* in the strain.

### Implementation of CICI

For steady-state images, we grew the log-phase cells in glucose, added rapamycin (Sigma catalog # 53123-88-9) to a final concentration of 1 ng/ul or equal volume of DMSO, and incubated for 1 hr. We then transferred the cells onto an agarose pad containing the same concentration of rapamycin / DMSO and further incubated for ∼10 mins for the cells to adjust to the solid medium. The agarose pad was then laid onto a glass coverslip and imaged under a Leica fluorescent microscope as described previously ^47^. Nine z-stacks were taken with an exposure time of 0.2 s and light intensity of 100% (SOLA SE light source) for both GFP and mCherry channels. For CICI time course, we directly placed log-phase cells onto an agarose pad with 1 ng/ul of rapamycin and started imaging immediately. The incubation time for each set of z-stack images was calculated as the duration between rapamycin addition and time of image acquisition.

### Image analysis

We developed Matlab software to analyze the z-stack fluorescent images ^29^. We first used the phase image to annotate the cell boundaries, and a composite image was generated from nine z-stack images where each pixel within the cell boundaries takes on the maximum intensity of this pixel among all the nine stacks. We then detected the locations of the green and red chromosome dots with intensities above a threshold. We recorded their x and y positions and then calculated the distance between the two dots. For cells containing more than one green or red dot, which might be due to DNA replication, we calculated the pairwise distances between each green and red dot and recorded the minimum distance. Green and red dots within 0.4 µm were considered as co-localized.

### CICI formation data analysis

The observed CICI formation time should include both the time for rapamycin to diffuse into the nucleus and the time for the two chromosomal loci to find each other. Assuming these are consecutive and independent Poisson processes, each of them following an exponential probability density function, we fitted the CICI formation curves with the following equation (the convolution of the two Poisson distributions):

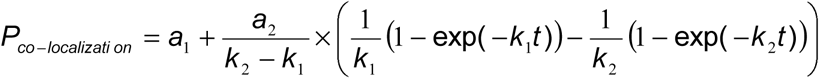

where *k*_1_ and *k*_2_ represent the rate of the two processes, and *a*_1_ and *a*_2_ determine the initial and final co-localization probability. We fixed the *k*_2_ (the rate for rapamycin to enter nucleus) to be 0.2 min^-1^ for all the CICI strains. *t*_*1/2*_ was calculated as ln2/*k*_1_. CICI efficiency is defined as the net gain of co-localization probability after reaching steady-state, which is calculated as:

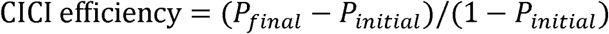

### *MAT* repair assay

We adapted the protocol developed by the Haber lab ^48^. Single colonies were grown overnight in 5 mL YEPD (YEP+ 2% Glucose, for no *HO* control) or YEPR (YEP+ 3% Raffinose, for HO induction) media with a yeast hormone “a factor” (10 ng/ul) at 30 °C to log phase. We noticed a high background switching frequency from a cell to α cell in YEPD where *GAL10pr*-*HO* was not induced, and 80 uM of “a-factor” (Zymo Research catalog # Y1004-500) was added to inhibit the growth of switched α cells before *HO* induction. We added 1 ng/ul rapamycin to these cultures to induce CICI formation between *MAT* and *HMR* for 1 hr (DMSO added as the control). We washed away the a-factor, added back rapamycin (or DMSO), and induced the expression of *GAL10pr*-*HO* for 1 hr with YEPG (YEP+ 3% Galactose). Glucose was then added to a final concentration of 2% to stop the *HO* expression, and cells were incubated for additional 1 hr to allow DNA repair. We next extracted genomic DNA from these samples and tested the genotype by PCR. To differentiate *MATa* vs. *MATα* (total switching), we used a forward primer which is immediately adjacent to the *MAT* locus and a reverse primer specific to either a or α sequence. To test the usage of *HML* vs. *HMR* among the switched population, we digested the *MATα* PCR products with BamHI and quantified the digested fraction using ImageJ. The undigested band (*MATα*) represents *the HML* usage, and two digested bands (*MATα-B*) represent the *HMR* usage. The total usage of *HMR* is calculated as the *MATα-B* band intensity divided by the total intensity of *MATa* and *MATα-B.*

## Acknowledgements

We thank Dr. Susan Gasser and Dr. James Haber for providing plasmids and yeast strains. We acknowledge all members in Bai lab for insightful comments on the manuscript. We also want to thank the members of the Center of Eukaryotic Gene Regulation at PSU for discussions and technical support. This work is supported by the National Institutes of Health (R01 GM118682).

## Author Contribution

L.B. and M.D. designed the experiments; M.D. performed most of the experiments and data analysis; F.Z. and Y.Y. helped with the experiments; M.D. and L.B. wrote the manuscript.

## Conflict of Interest Statement

The authors declare no conflict of interest.

